# Immune Checkpoint B7-H3 Inhibition Reprograms Acute Neuroinflammation and Protects the Brain After Acute Ischemic Stroke

**DOI:** 10.64898/2026.02.26.708106

**Authors:** Arun Kumar Boda, Kit-Kay Mak, Weiguo Li, Bharath Chelluboina

**Author notes:** Corresponding Author: Bharath Chelluboina, Department of Pharmacy Practice, University of Illinois, Chicago, IL, 60612.

## Abstract

**BACKGROUND AND PURPOSE:** The role of immune checkpoint B7-H3 in acute ischemic stroke prognosis and post-stroke immunosuppression remains uninvestigated, despite the clinical significance of immune checkpoints with inflammaging and post-stroke infections. We recently reported the neuroprotective effects of acute B7-H3 inhibition after stroke in adult and aged mice. In this study, we investigated the mechnsistic association of regulating the cerebral induction of B7-H3 after acute ischemic stroke on brain damage, neuroinflammation, vascular integrity, host defense gene regulation, and functional outcomes.

**METHODS:** C57BL/6 mice were subjected to transient middle cerebral artery occlusion and injected (i.v.) with either B7-H3 siRNA or a negative (non-targeting) siRNA at 5 min after reperfusion. At 24 hours of reperfusion, magnetic resonance imaging (MRI) of the mouse brain was performed using a 9.4 T scanner to assess brain damage (T2, ADC, and Kurtosis). Real-time qPCR and NanoString nCounter® Neuroinflammation Panels were used to determine the acute changes in overall neuroinflammatory functions that are mediated by B7-H3. Motor function (beam walk, rotarod tests, and grip strength) was assessed between days 1 and 7 of reperfusion.

**RESULTS:** Early inhibition of B7-H3 after stroke significantly reduced the brain damage and promoted the functional outcomes. Post-stroke neuroinflammation was reprogrammed with B7-H3 inhibition towards balancing of neuroprotective anti-inflammatory mechanisms without compromising the immune response that is crucial for preventing post-stroke infections.

**CONCLUSIONS:** Our results demonstrate that the induction of B7-H3 during the acute period after stroke is a mediator of post-stroke neuroinflammation and secondary brain damage.

## Introduction

Immune checkpoints dysregulate in the aging brain and in cerebrovascular diseases.^1^ Immune checkpoints shape the balance between neuroprotection and injury, but their therapeutic modulation is complicated by infection risk and timing constraints after stroke.^1,2^ CD276 (B7-H3) acts as a bidirectional immune checkpoint that modulates T□cell activity, glial neuroinflammation, neurovascular integrity, and shapes injury/repair mechanisms in CNS diseases.^3,4^ CNS infection and meningitis show that B7-H3 upregulation exacerbates BBB disruption, cytokine storms, and neurological injury.^5,6^ Further, B7-H3 knockout mice have shown significantly reduced neuroinflammation and protection after experimental autoimmune encephalomyelitis.^7^ Despite the temporal profile of B7□H3 expression in experimental stroke aligning with the early wave of neuroinflammation, no studies have yet investigated its modulation after acute ischemic stroke or its functional significance in inflammatory signaling. Therefore, to fill that void, we recently determined the effect of early knockdown of B7-H3 after stroke in adult aged mice and reported our preliminary outcomes at Neuroscience 2025^8^ and International Stroke conference 2026^9^, We extended our studies to mechanistically determine B7-H3 role in neuroprotection and host defense by determining the impact of B7-H3 knockdown on post-stroke brain damage and motor function recovery in adult mice.

## Materials and Methods

All animal procedures were approved by the University of Illinois Chicago Research Animal Resources and Care Committee. Animals were cared for in compliance with the Guide for the Care and Use of Laboratory Animals [U.S. Department of Health and Human Services publication no. 86–23 (revised)]. All procedures were conducted in compliance with the “Animal Research: Reporting of In Vivo Experiments” guidelines. Mice were randomly assigned to experimental groups. Mice with no noticeable neurological deficits or no sign of infarction in MRI scans on day 1 of reperfusion, and those that showed a hemorrhage upon euthanasia were excluded.

Adult C57BL/6 mice of both sexes (3 months) and aged C57BL/6 males (18 and 24 months) (Jackson Laboratories USA) were used in the study. Mice were subjected to 1h (adult) of transient middle cerebral artery occlusion (MCAO) under isoflurane anesthesia (Somniflow) followed by B7-H3 siRNA cocktail administration at 5 min of reperfusion. At 1 day of reperfusion, all mice underwent Magnetic Resonance imaging (MRI) to determine brain damage, cerebrovascular mechanisms assessed (qPCR, Immunoblot, and nCounter Pathway focused Assays). Functional outcomes were assessed for 7 days of reperfusion. All procedures except MRI (ADC and Kurtosis) and Nanostring nCounter pathway-based assays were demonstrated in our earlier studies.^10,11^ The detailed methods were provided in the supplemental methods.

## Results

### 1. B7-H3 inhibition reduced brain damage and promoted functional outcomes after acute ischemic stroke

B7-H3 expression in the brain increased with aging, and after stroke, it increased as early as 6h in males and 12h in females (Supplemental Fig 1). To evaluate the neuroprotective effect of B7-H3 inhibition in ischemic stroke, we determined the effect of B7-H3 siRNA treatment immediately after stroke on brain damage at 24h of treatment. The B7-H3 siRNA-treated mice showed significantly reduced infarct volume compared with the Neg siRNA-treated mice (Fig. 1A). The reduction of tissue injury indicates that B7-H3 knockdown attenuates early ischemic neuronal damage. To characterize the structural and physiological changes induced by injury, we performed T2-weighted and diffusion-weighted MRI and generated T2, Apparent Diffusion Coefficient (ADC), and diffusion Kurtosis(K) maps. Diffusion-weighted MRI is sensitive to microstructural changes in tissue caused by early cytotoxic edema following arterial occlusion and typically shows marked reductions in ADC in ischemic tissue and an increase in diffusion kurtosis due to highly restricted water diffusion. As demonstrated by the derived metric maps and histogram analysis, we found that, when compared with the B7-H3 siRNA-treated mice, the ipsilateral hemisphere of Neg siRNA-treated mice showed consistent high T2 (Fig. 1C), decreased ADC (Fig. 1D), and increased Kurtosis values (Fig. E) in all three affected brain areas (cortex, striatum and hippocampus). These structural changes indicate ameliorated cytotoxic edema and reduced infarct progression after treatment with B7-H3 siRNA. Early neurological recovery was assessed using a battery of motor performance tests. Post-stroke motor function recovery (rotarod test and beam walk test) was significantly higher in the B7-H3 siRNA cohorts between days 1-7 of reperfusion compared with the age-matched Neg siRNA cohorts (n = 7/group for *p < 0.05) (Fig. 1B). However, grip strength did not improve significantly, although it showed a positive trend (Supplementary Fig 3). These improvements indicate that B7-H3 blockade confers functional benefits during the subacute phase of recovery.

**Figure 1.**
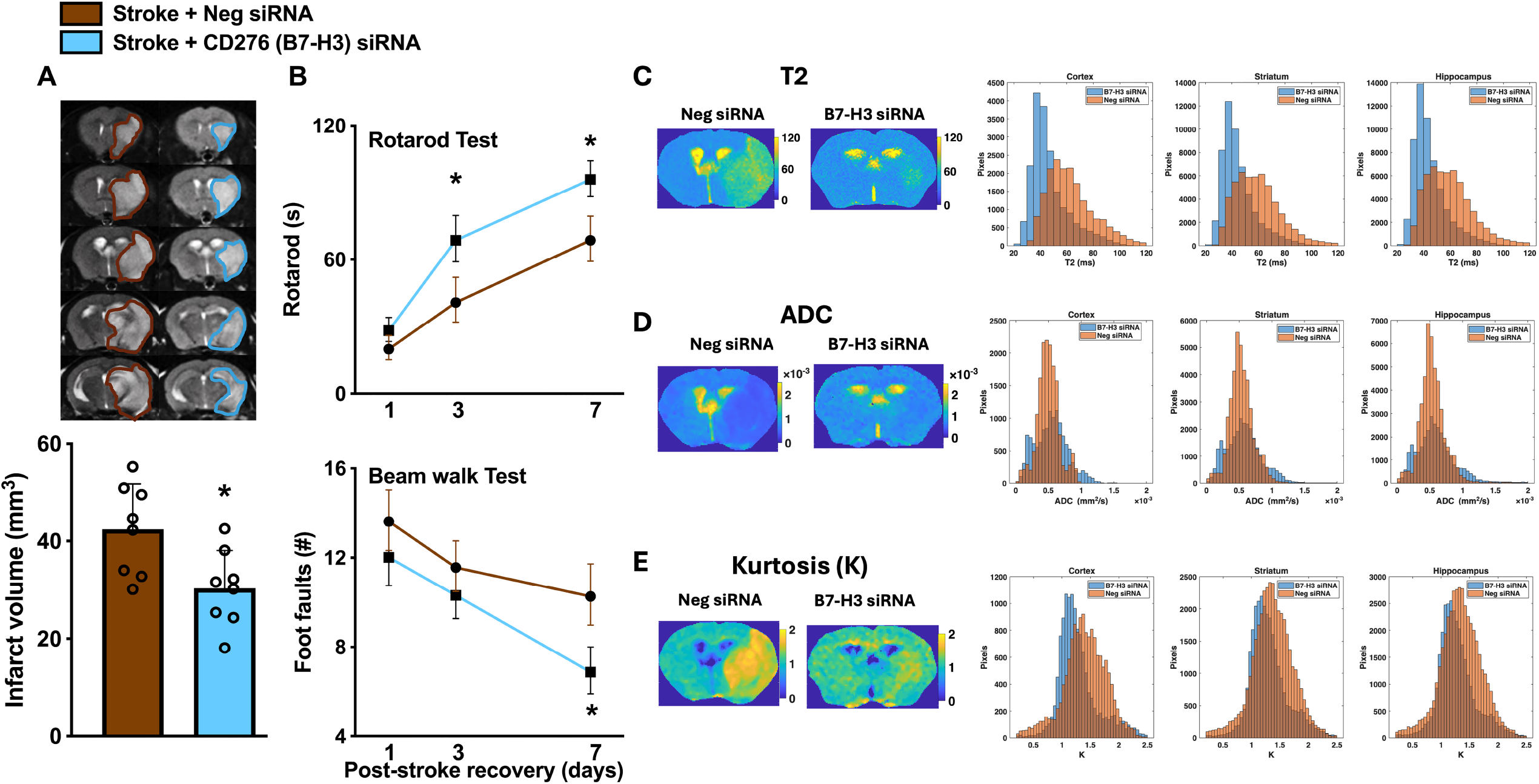
MRI mapping and functional analysis of brain damage and functional outcomes following MCAO in mice. (A) Representative MRI showing infarcted brain damage 24h after MCAO/R in Neg-siRNA and B7-H3 siRNA-treated mice. Quantitation of infarct volume measured based on MRI images. Mann–Whitney U post□test (N = 8-10, P < 0.05). (B) Motor functions, including Beam Walk performance, were measured over 7 days of reperfusion and the Rotarod test, representing latency to fall, was assessed over 1 week to determine the impact of early inhibition of B7-H3. A non-parametric two-way repeated measures ANOVA with Sidak’s multiple comparison test. (N=8-10, P < 0.05). (C) T2 maps and T2 histograms of three brain areas (Cortex, striatum, and hippocampus) representing edema and lesion. Unit of T2: millisecond. (D) Apparent diffusion coefficient (ADC) maps and ADC histograms acquired at 24 h of post-treatment of MCAO/R, representing the microstructural changes in the lesion areas. Unit of ADC: mm^2^/s. (E) Diffusion kurtosis (K) maps and K histograms acquired at 24 h post-treatment in MCAO/R, reflecting the inherent heterogeneity of microstructural architecture changes in the lesion.

### 2. Post-stroke B7-H3 is proinflammatory and affects BBB integrity

To determine the B7-H3 regulated inflammatory response, we quantified the mRNA expression of six critical inflammatory mediators implicated in stroke pathogenesis [toll-like receptor 4 (TLR-4), interleukin-1 beta (IL-1β), matrix metalloproteinase-9 (MMP9), vascular endothelial growth factor (VEGF), CC-chemokine ligand 3 (CCL3/MIP-1α), and tumor necrosis factor-alpha (TNF-α)]. All six inflammatory genes showed significant upregulation after stroke, demonstrating robust neuroinflammatory activation at 24h of stroke. TLR-4, IL-1β, MMP9, and VEGF expression in the peri infarct zone was significantly reduced by B7-H3 knockdown after acute stroke indicate B7-H3 regulates upstream pattern recognition receptor (TLR-4) that drives NF-κB leads to attenuation of the TLR4-NF-κB signaling cascade that amplifies inflammatory injury and oxidative stress, substantial dampening of IL-1β-driven pathological processes including endothelial activation, neutrophil recruitment, and vascular permeability (Figure 2 A, B, D & E). By reducing MMP9 at 24h, B7-H3 knockdown preserves BBB structural integrity and reduces the risk of hemorrhagic transformation. Further, VEGF regulated by B7-H3 knockdown at 24h reduces pathological vascular permeability. Interestingly, in contrast to the coordinate reduction of TLR4, IL-1β, MMP9, and VEGF, two critical immune genes (TNF-α and CCL3) remained unchanged after B7-H3 knockdown (Figure 2C & F). This indicates B7-H3 inhibition preserves CCL3 ability to recruit immune cells in response to bacterial infections, a critical mechanism to prevent stroke-induced immunosuppression.

**Figure 2.**
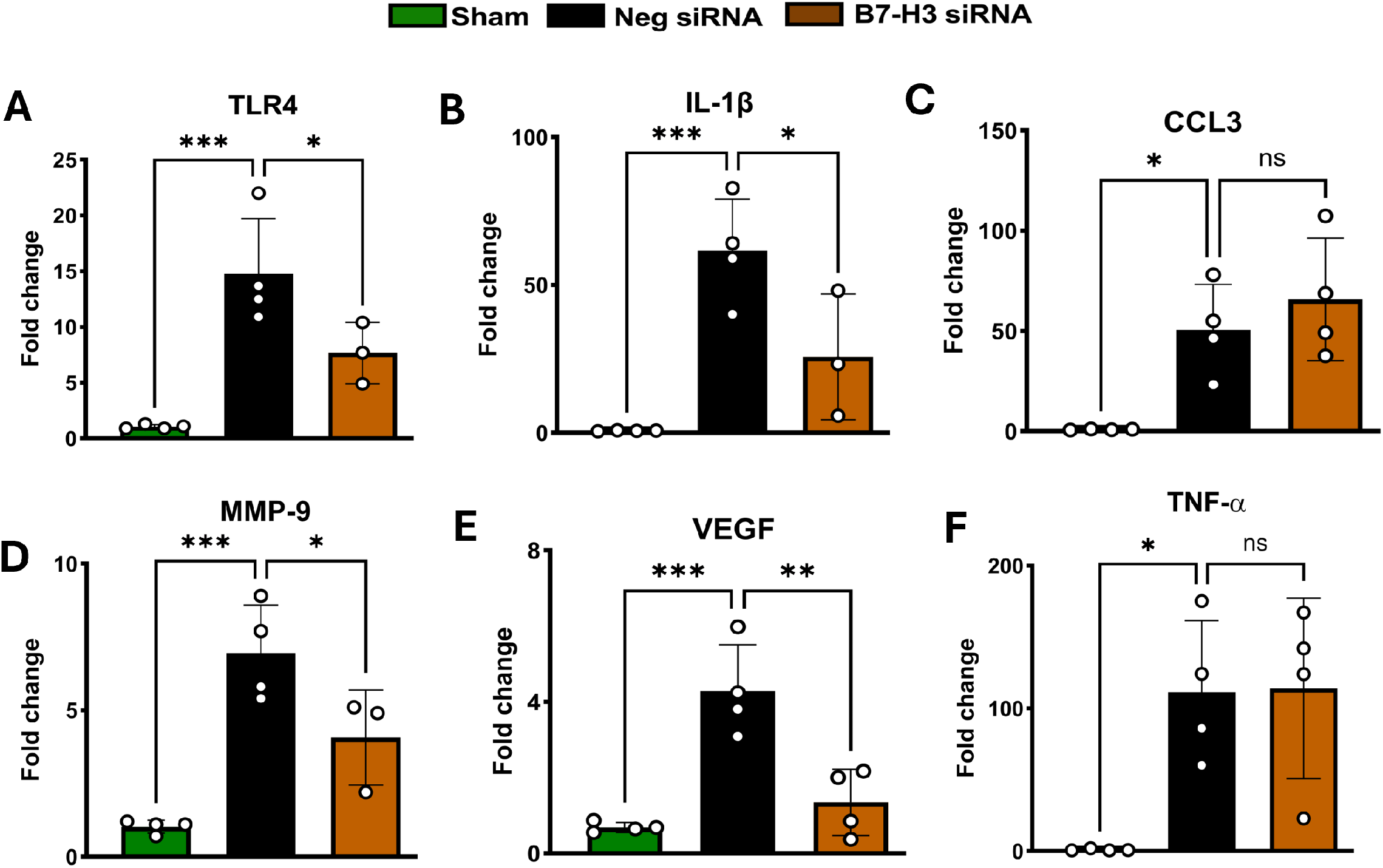
Proinflammatory mediators’ expression following post-treatment of Neg siRNA and B7-H3 siRNA at 24 h.(A-F) Quantitative mRNA expression analysis of proinflammatory mediators including TLR4, IL-1β, CLL3, MMP9, VEGF and TNF-⍰in penumbra tissue at 24 h of MCAO/R. Statistical analysis was performed using One-way ANOVA test (N = 4-5 per group).(G) RNA seq heat map depicting gene expression profile of inflammatory signaling pathways at 24 h of post-treatment.

We also evaluated the stroke-induced modulation of multiple immune checkpoints, including PD-1, PD-L1, CTLA4, and TIM-3 in the B7-H3 inhibition condition. Knockdown of B7-H3 did not alter the expression of any other immune checkpoints, indicating the absence of cross-regulation. These findings suggest that the B7-H3 pathway operates through a distinct inflammatory response during acute ischemic stroke (Supplementary Fig 2).

### 3. B7-H3 knockdown promotes post-stroke inflammatory signaling toward tissue protection without potential immunosuppression

The neuropathology panel captured pathways related to tissue remodeling, immune regulation, inflammatory resolution, and host defense in stroke mice that underwent B7-H3 siRNA and Neg siRNA treatments. Several inflammation-associated genes linked to tissue repair, extracellular matrix stabilization, and regulated innate immune responses were robustly upregulated in stroke mice treated with B7-H3 siRNA compared with Neg siRNA controls. In parallel, anti-inflammatory, inhibitory immune signaling, and immune-regulatory genes were further elevated in the B7-H3 siRNA group suggests enhanced immune restraint and promotion of inflammatory resolution (Figure 3A-C). In addition, innate immune sensing and response pathways remained preserved following B7-H3 knockdown indicating that B7-H3 knockdown does not suppress core host defense mechanisms despite attenuating potentially damaging inflammatory outputs. Genes associated with immune cell migration and endothelial interaction, including Jam2 and Chn2, remained suppressed in both stroke groups and showed no induction following B7-H3 knockdown suggest enhanced immune restraint and reduced leukocyte trafficking following acute B7-H3 knockdown. Additionally, immune differentiation and cellular stress responses (including Eomes, Ugt8a, Nwd1, Fgf13, Akt2, Spib, Cryba4, Cpa3, and Rad51b) remained downregulated in both groups suggest B7-H3 knockdown does not broadly reverse stroke-associated immune suppression (Figure 3B).

**Figure 3.**
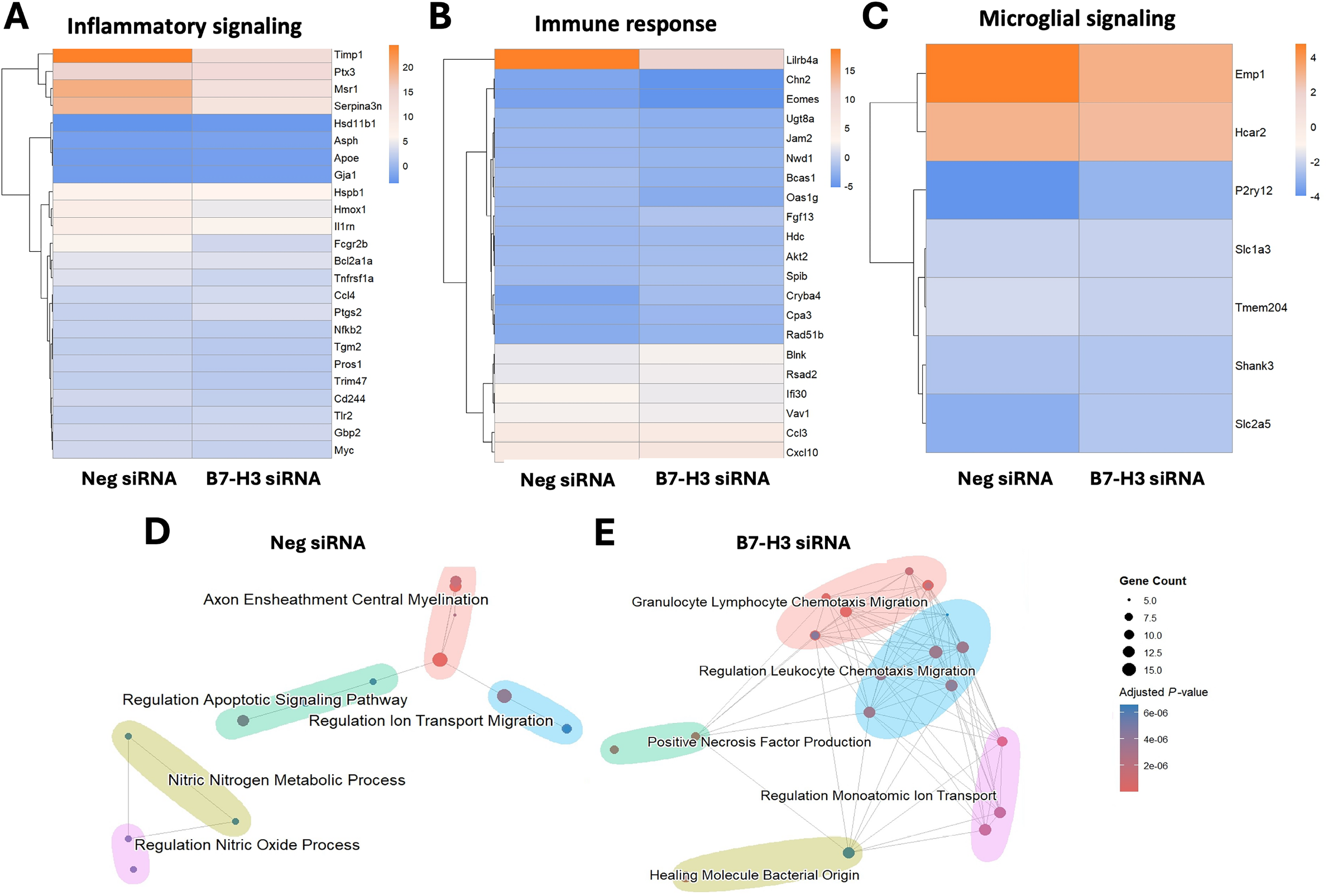
Gene expression profile shows neuroinflammation, immune signaling and enrichment maps of MCAO operated mice brain samples at 24h. RNA-seq heatmap shows (A) immune response, and (B) microglial signaling associated genes following ischemic stroke and B7-H3 mediated mechanisms. (C & D) Enrichment maps show functional significance of post-stroke B7-H3 inhibition (N=3).

Enrichment map (emap) analysis reveals distinct immune and tissue-response programs after stroke with B7-H3 knockdown. Enriched Gene Ontology (GO) biological processes were visualized as network-based maps, where node size reflects gene count and edge connectivity indicates shared genes between pathways.^12^ In Neg siRNA stroke mice, enriched pathways were primarily associated with neuronal structural integrity and intrinsic stress responses, including axon ensheathment and central myelination, regulation of apoptotic signaling, ion transport, and nitric oxide-related metabolic processes suggests a compartmentalized response dominated by neuronal injury and metabolic stress with minimal immune coordination, consistent with post-stroke immune dysfunction (Figure 3D). In contrast, B7-H3 siRNA-treated mice showed a distinct enrichment network characterized by dense, highly interconnected immune-related clusters that indicate coordinated immune activation(Figure 3E).^13^ However, these immune pathways also showed shared gene programs and B7-H3 knockdown shifts post-stroke responses from neuronal stress towards organized immune regulatory and host-defense mechanisms.

## Discussion

In this study, we demonstrate that early inhibition of immune checkpoint B7-H3 after ischemic stroke offers neuroprotection by selectively regulating inflammatory pathways associated with BBB disruption in acute ischemic stroke and host response while operating independently of classical T cell immune checkpoints. These findings suggest a functional role of cerebral B7-H3 within the neuroinflammatory network rather than a broad immunosuppressive cascade.

Although B7-H3 knockdown does not inhibit other immune checkpoints, it exerts selective regulation of inflammatory mediators (TLR-4, IL-1β, MMP9, and VEGF) that drive BBB and secondary brain damage. This suggests that B7-H3, by regulating the inflammatory network during acute stroke, coordinates the expression of functional genes involved in BBB disruption while independently maintaining immune surveillance by preserving CCL3 and TNF-α.

Significant reduction in cerebral TLR-4 expression with B7-H3 knockdown after acute stroke suggests that B7-H3 and TLR-4 may be co-regulated through shared cellular/transcriptional mechanisms, and that B7-H3 knockdown disrupts this signaling axis, likely through anti-inflammatory effects that attenuate the downstream NF-κB-driven transcriptional cascade. The significant reduction of IL-1β expression following B7-H3 knockdown further supports the downstream consequence of attenuated TLR4-NF-κB signaling.^14^ The reduction of both TLR-4 and IL-1β by B7-H3 knockdown creates a synergistic anti-inflammatory effect.^14^ MMP9 reduction with B7-H3 knockdown shows a direct BBB protective mechanism at the acute phase of stroke.^15^ Interestingly, reduction of MMP9 while preserving CCL3 and TNF-α with B7-H3 knockdown is therapeutically advantageous, as neutrophils recruited by CCL3 are a primary source of MMP9 in the peri-infarct region.^16^ Thus, B7-H3 knockdown drives inflammatory reprogramming by preserving CCL3-mediated neutrophil recruitment that maintains antimicrobial surveillance capacity while reducing MMP9 expression that limits proteolytic BBB damage.^17^ Furthermore, early VEGF inhibition (0-24 h) reduces infarct volume by 20-30%, attenuates BBB disruption, preserves tight junction proteins, and reduces MMP9 expression.^18^ B7-H3 knockdown reduced acute-phase VEGF expression that exerts predominantly harmful effects on BBB integrity and edema formation. However, caution is needed as VEGF at subacute timepoints (3-7 days) is crucial for angiogenesis and recovery.^19^ The preservation of CCL3 and TNF-α expression despite B7-H3 knockdown provides mechanistic insight into B7-H3’s selective regulatory functions. While excessive CCL3-mediated inflammation can exacerbate ischemic injury, CCL3 is essential for antimicrobial defense, immune cell trafficking to infection sites, and coordination of cellular responses to bacterial invasion.^20^ CCL3-deficient mice show increased susceptibility to bacterial and viral infections, demonstrating its critical role in host defense.^21^ The preservation of CCL3 expression following B7-H3 knockdown ensures stroke patients retain the capacity to recruit immune cells in response to bacterial infections, which is critically important given that stroke-induced immunosuppression dramatically increases susceptibility to pneumonia and urinary tract infections. Therefore, B7-H3 knockdown may enable beneficial immune surveillance while preventing excessive vascular destruction. Furthermore, preserved TNF-α and reduced IL-1β, after B7-H3 inhibition, are therapeutically advantageous as it promotes tissue repair, immune regulation, and host defense while reducing acute neurotoxicity.

These findings demonstrate that acute B7-H3 knockdown following ischemic stroke induces a selective reprogramming of immune and inflammatory responses characterized by enhanced inhibitory myeloid signaling, reduced leukocyte trafficking, and increased expression of tissue-protective and immune-regulatory mediators, thereby representing a promising strategy for mitigating post-stroke complications. We limit our study to the acute phase of ischemic stroke, as immunomodulation at this phase has dual effects of neuroprotection and host defense.

## Abbreviations

B7-H3: also known as CD276
MRI: magnetic resonance imaging
MCAO: middle cerebral artery occlusion
siRNA: small interfering RNA
GAPDH: glyceraldehyde 3-phosphate dehydrogenase.

## Author Contributions

B. C conceived and designed the study. A.K.B performed experiments. B.C and W. L performed MRI analysis. MKK performed gene expression analysis. B.C, A.K.B, and K.K.M wrote, revised and finalized the manuscript. All authors contributed to the article and approved the submitted version.

