## Supplementary material for "Immune Checkpoint B7-H3 Inhibition Reprograms Acute Neuroinflammation and Protects the Brain After Acute Ischemic Stroke": n/a

**Abbreviated title**: B7-H3 induction drives acute ischemic stroke pathogenesis

**Materials and methods:**

All animal procedures were approved by the University of Illinois Chicago Research Animal Resources and Care Committee. *Animals were cared for in compliance with the Guide for the Care and Use of Laboratory Animals [U.S. Department of Health and Human Services publication no. 86–23 (revised)].* All procedures were conducted in compliance with the "*Animal Research: Reporting of In Vivo Experiments*" guidelines.[1] Mice were randomly assigned to experimental groups. Mice with no noticeable neurological deficits or no sign of infarction in MRI scans on day 1 of reperfusion, and those that showed a hemorrhage upon euthanasia were excluded.

**Focal cerebral ischemia**

Adult C57BL/6 mice of both sexes (12 weeks old) (Jackson Laboratories USA) were subjected to 1h (adult) of transient middle cerebral artery occlusion (MCAO) under isoflurane anesthesia (Somniflow) followed by 1 to 7 days of reperfusion as described earlier.[2-5] Sham-operated animals served as the control. Regional cerebral blood flow (rCBF) and physiological parameters (pH, $PaO_2$, $PaCO_2$, hemoglobin, and glucose) were monitored, and rectal temperature was maintained at 37 ± 0.5°C during surgery and recovery from anesthesia. Post-surgical care and monitoring were provided as specified in the IMPROVE guidelines.[6]

**siRNA treatment**

Mice were injected via a retroorbital route with 25 nmol (100 µl) of Ambion® silencer select in vivo grade CD276 siRNA or negative siRNA (Neg siRNA; cat# 4404020; siRNA ID#: s813) (a cocktail of 3 siRNAs in each case; Thermo-Fisher USA) mixed with 50 µl polyethylene glycol-liposome in vivo transfection reagent and 10 µl transfection enhancer (Altogen Biosystems USA). The sequences (5'->3') of the CD276 siRNAs used as a cocktail are (cat#4457308 siRNA ID: s97988 sense seq ACCCUACGCUGCUCCUCCUUUtt, antisense seq

AAACCAGCAGCGUAGGGUgg; siRNA ID: s97987 sense seq CACCCAGCCUUACUUUUtt antisense AAAAGUAAGGUGUGGGUGgt; siRNA ID: s97986 sense seq CCUGCUUUGUGAGAGCAUCCAtt antisense UGGAUGCUCACAAAGCAGGtg (Thermo-Fisher USA). The siRNA cocktails were injected at 5min of reperfusion.

**Magnetic resonance imaging (MRI)**

At day 1 of reperfusion following transient MCAO, mice were anesthetized with isoflurane and subjected to MRI. Respiration rate was monitored during the imaging. None of the animals showed any distress or mortality during or after scanning. MRI scans were analyzed by a person blinded to the study groups to estimate infarct volume (T2-MRI). Briefly, Mouse brain MRI was performed using a 9.4 T scanner with a gradient coil of 100 G/cm (Agilent) and a 39 mm ID RF coil (Rapid). A multiple slice multiple echo Carr-Purcell-Meiboom-Gill sequence was used for measuring transverse relaxation times (T2) with acquisition parameters: TR = 4 s, 12 TEs with the first TE = 6.1 ms and the echo space = 6.1 ms, slice thickness = 1 mm, FOV = 19.2 × 19.2 mm, matrix = 128 × 128, NEX = 4. A fast spin echo sequence with diffusion gradient on slice-selection direction was applied with following acquisition parameters: TR = 2 s, effective TE = 22 ms, echo train length = 8, slice thickness =1.0 mm, FOV = 19.2 × 19.2 mm, matrix = 128 × 128, NEX = 4, $\delta/\Delta = 5/10$ ms, 8 b-values: 0, 650, 800, 1000, 1250, 1500, 2000, 2500 s/mm2.

Data Postprocessing: All data were processed using MATLAB (MathWorks). The T2 and diffusion parameter maps were generated pixel-wise using the monoexponential model. To characterize non-Gaussian water diffusion arising from post-stroke tissue microstructural complexity, the diffusion kurtosis imaging (DKI) framework was applied. In this model, the normalized signal attenuation is expressed as a second-order Taylor expansion of the logarithm of the signal with respect to b-value, yielding estimates of both diffusion and kurtosis.[7]

**Motor Function Analysis**

Motor function was evaluated by the rotarod test (4 min on a cylinder rotating at 15 rpm) and beam walk test (number of foot faults while crossing a 120-cm long beam) at days 1 and 7 of reperfusion by a person blinded to the groups, as described earlier. Mice were trained in the tasks for 3 days before MCAO. In addition, we used grip strength assessment to further evaluate the functional regain. Briefly, before MCAO surgery, mice were allowed to practice walking on a rotarod (Panlab Harvard apparatus) at a speed of 5 rpm, 10 rpm, and 15 rpm (revolutions per minute) for three days. After surgery stress, we evaluated the ability of the mice in each group to walk on the rotarod, which was accelerated at 15 rpm for 4 min. The time at which each mouse fell off the rod was recorded as the retention time, and the data was subjected to statistical analysis. Muscle strength was measured by a grip strength meter using a digital force gauge (BIO-GS4, BIOSEB in vivo research instruments) with mesh pull bars. The peak strength applied to the gauge was recorded in five, and the mean value was calculated. The mouse forelimb strength was measured by positioning the gauge horizontally. Both front paws were placed on the pull bar, and the mouse gently pulled in a straight direction away from the bar until releasing it.

**Real-time PCR and Western Blotting**

RNA isolation and qPCR:

The tissue was homogenized and total RNA was isolated using Purelink RNA kit (cat#12183025 Invitrogen). The RNA isolation was done according to the kit manufacturer's protocol. The purified RNA was reversely transcribed by using cDNA synthesis kit (cat# 4387406 Applied Biosciences). The PCR amplicon was carried out using SyBr green mix (cat# 4309155 Applied Biosciences) with the following primers referred in supplementary table 1. Quantitative real-time

PCR was performed using Quantstudio Flex 3/5 system (Thermo Scientific, USA). 2−ΔΔCt method was used to calculate the relative expression levels of target genes.

**Supplementary Table 1: Real-time PCR Primers**

| **Gene** | **Sequence (5' to 3')** |
|---|---|
| B7-H3 | CTACAGCTGCCTGGTACGCAA |
| | CAGAGGGTTTCAAGAGGCCGTA |
| TIM-3 | CCCTGCAGTTACACTCTACC |
| | GTATCCTGCAGCAGTAGGTC |
| LAG-3 | CTCCATCACGTACAACCTCAAGG |
| | GGAGTCCACTTGGCAATGAGCA |
| MMP9 | GACATAGACGGCATCCAGTATC |
| | GTGGGAGGTATAGTGGGACA |
| TLR4 | AAATGCACTGAGCTTTAGTGGT |
| | TGGCACTCATAATGATGGCAC |
| TNF-α | CAGGCGGTGCCTATGTCTC |
| | CGATCACCCCGAAGTTCAGTAG |
| VEGF | TGGTTCTTCACTCCCTCAAATC |
| | GGTCTCTCTCTCTCTTCCTTGA |
| IL-1β | GCAACTGTTCCTGAACTCAACT |
| | ATCTTTTGGGGTCCGTCAACT |
| PD-L1 | GGCAGGAGAGGAGGACCTTA |
| | TGCAGCTTGACGTCTGTGAT |

| | |
|---|---|
| CTLA4 | GCCAGTGGTTCCAAAGGTTG |
| | CACTGTGGGACGACACTGAT |
| 18S | CGCCGCTAGAGGTGAAATTCT |
| | CGAACCTCCGACTTTCGTTCT |

**Immunoblotting**

The protein lysis buffer contains RIPA buffer (cat#89900 Thermo Scientific) with protease and phosphatase inhibitor cocktail (cat#78441, Thermo scientific, 100X). Protein samples were subjected to Bolt 4-12% Bis-Tris plus wedge well gel and transferred to nitrocellulose membranes. Membranes were blocked and incubated with primary antibodies (B7-H3 cat#AB134161, Abcam (knockout validated) GAPDH cat# sc-47724, Santa cruz) overnight and secondary antibodies were conjugated with HRP. After washing five times with 1X PBST, the protein bands were visualized using an electrogenerated chemiluminescent (ECL) solution and Azure Biosystem-500.

**The NanoString nCounter® neuroinflammation pathways analysis**

The gene expression profiling was performed using the NanoString nCounter® Neuroinflammation Panel (NanoString Technologies, USA). This panel targets a curated set of 770 genes involved in neuroinflammatory, immune, and injury-related pathways. RNA was isolated from brain penumbra tissue from mice subjected to MCAO and treated with either negative control siRNA (Neg siRNA) or B7-H3-targeting siRNA (B7-H3 siRNA). RNA quality and quantity were assessed prior to hybridization according to the manufacturer's instructions.

Raw NanoString counts were processed and normalized using nSolver™ Analysis Software (NanoString Technologies). Background correction was applied using built-in negative control probes, followed by normalization to internal positive controls and selected housekeeping genes to account for technical variability. Differential gene expression analysis was performed within nSolver by comparing B7-H3 siRNA treated mice against Neg siRNA stroke controls with sham mice as the reference group. The RNA samples consisted of three biological replicates (n = 3 per group). Genes were considered significantly differentially expressed if they met the following criteria: adjusted p-value $\leq 0.05$ and fold change (FC) $\geq 2$ (FC $\geq 2$ or FC $\leq -2$). Genes passing these thresholds were further curated based on biological relevance to inflammatory signaling, immune regulation, and host defense pathways. Heatmaps were generated using RStudio (R version 4.5.1) with appropriate visualization packages (pheatmap, ggplot2). Expression values were $\log_2$-transformed and scaled by gene to facilitate comparative visualization across samples. Hierarchical clustering was performed on genes using Euclidean distance and complete linkage to identify patterns of co-regulated gene expression. Color gradients represent relative expression levels, with warmer colors indicating higher expression and cooler colors indicating lower expression.

**Gene Ontology enrichment and enrichment map analysis**

Gene Ontology (GO) enrichment analysis was performed to identify biological processes associated with differential gene expression following B7-H3 knockdown after MCAO. Gene symbols were converted to Entrez Gene IDs using the ‘bitr’ function from the ‘clusterProfiler’ package to ensure compatibility with downstream enrichment analyses. Mouse gene annotation was obtained from the org.Mm.eg.db database.

GO enrichment analysis was conducted using the enrichGO function implemented in the clusterProfiler package with a focus on the Biological Process (BP) ontology. The analysis was performed using Entrez Gene IDs as input, with the Benjamini-Hochberg (BH) method applied for multiple testing correction. Enriched GO terms were considered significant at a p-value cutoff of

0.05 and a q-value cutoff of 0.2. For improved interpretability, enriched gene identifiers were converted back to gene symbols.

Enrichment map (emap) visualizations were generated using the 'enrichplot' package within RStudio. In the emap plots, each node represents an enriched GO biological process, with node size proportional to the number of genes associated with the term. Edges indicate gene overlap between GO terms, reflecting shared biological pathways and coordinated functional relationships. Node color corresponds to adjusted p-values, with increased color intensity indicating higher statistical significance. All analyses and visualizations were performed using RStudio (R version 4.5.1), with required packages installed via BiocManager.

**Statistical Analysis**

All statistical analyses were performed using standard parametric or non-parametric approaches. Two groups were compared with the Mann-Whitney U test and multiple groups with one-way ANOVA with Tukey's multiple comparisons test. Data collected repeatedly from the same set of subjects at different time points (e.g., rotarod and beam-walk tests) were analyzed using a nonparametric two-way repeated-measures ANOVA with Sidak's multiple-comparisons test.

**Supplementary Data**

**Cerebral B7-H3 expression significantly increases with aging and acutely after ischemic stroke in mice and B7-H3 siRNA prevented upregulation of B7-H3 mRNA and protein induction after acute ischemic stroke.**

Basal levels of B7-H3 mRNA were measured in in age (3, 18, and 24 months) in CB57BL/6J male mice brain (supplementary fig. 1A) and were upregulated in aging. To determine B7-H3 temporal expression in the brain after acute ischemic stroke in male and female mice, brain tissues from adult male and female mice that were subjected to 60 min transient MCAO and 6, 12, and 24h of reperfusion were used to assess mRNA expression and determine time to target. B7-H3 mRNA levels in the brain were significantly increased within 24h of reperfusion. In adult male mice, transient MCAO led to a significant upregulation of B7-H3 mRNA as soon as 6h of stroke whereas 12h of stroke showed significant upregulation in adult female (n = 4/group, *p < 0.05) (supplementary fig. 1B). We used adult male mice to determine the effect of B7-H3 knockdown using ambion in vivo ready siRNA. B7-H3 siRNA treatment immediately after reperfusion led to significant knockdown of B7-H3 mRNA as well as protein compared to neg siRNA treatment assessed at 24h of reperfusion by approximately 50% compared to negative control siRNA-treated animals (n = 4-6/group, (Supplementary fig. 1C & D). This significant reduction in B7-H3 induction validates the efficacy and specificity of the selected dose, route, and timing of treatment with B7-H3 siRNA at the 24h post-stroke timepoint.

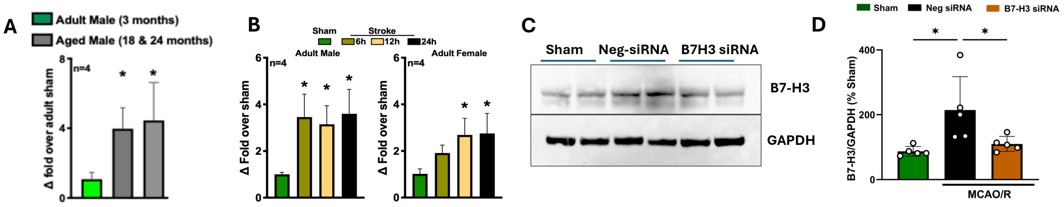


**Supplementary Fig 1:** (A) Relative B7-H3 mRNA expression in brain of adult mice (3 months) compared to aged mice (18 months and 24 months). (B) Time dependent B7-H3 mRNA expression following MCAO/R, showing early induction at 6h in male and 12h in female mice. (C) Represent western blot images depicting B7-H3 protein expression in penumbra tissue of Neg siRNA and B7-H3 siRNA post-treatment at 24h of MCAO/R. (D) Quantification of western blot intensity that show B7-H3 protein significantly reduced after stroke and B7-H3 knockdown. Data represents as mean ± SEM (N = 4-5 per group). One- Way ANOVA, followed by Dunnett's post hoc test.

**B7-H3 induces parallel to other immune checkpoints after stroke but drives independent mechanisms.**

Stroke induced upregulation of multiple immune checkpoints but B7-H3 inhibition is target specific without cross regulation of other immune checkpoints in acute stroke. Stroke triggered widespread upregulation of other immune checkpoints (PD-1, PD-L1, CTLA-4, TIM-3, and LAG-3) along with B7-H3 demonstrating the robust stroke responsive transcriptional activation. To determine whether B7-H3 functions as master regulator of immune checkpoints expression in acute stroke, peri infarct tissues from stroke with neg siRNA and B7-H3 siRNA treatments along with location matched sham were assessed at 24h of stroke. Despite a significant reduction of the target gene, B7-H3 knockdown did not significantly alter the expression of any other immune checkpoint molecules examined (Supplementary figure 2). Lack of cross regulation between the B7-H3 with other immune checkpoints indicate that these pathways are respond through distinct and parallel inflammatory responses at the acute phase of ischemic stroke.

CTLA-4 expression in B7-H3 siRNA-treated animals (~5 fold) showed no significant difference compared to Neg siRNA controls (~10 fold; $p > 0.05$). Similarly, LAG-3 expression remained unchanged between treatment groups (B7-H3 siRNA: ~2 fold vs. Neg siRNA: ~4 fold; $p > 0.05$). PD-L1 expression was statistically indistinguishable between the B7-H3 siRNA and Neg siRNA groups ($p > 0.05$). PD-1 and TIM-3 likewise showed no significant differences between B7-H3 siRNA and Neg siRNA treatment groups (both $p > 0.05$). For PD-1, expression levels were approximately 25-fold in Neg siRNA animals and 12.5-fold in B7-H3 siRNA animals, representing an apparent 50% reduction that did not achieve statistical significance. TIM-3 expression showed a similar pattern, with ~3-fold in Neg siRNA and ~1.5-fold in B7-H3 siRNA groups, again without reaching statistical significance.

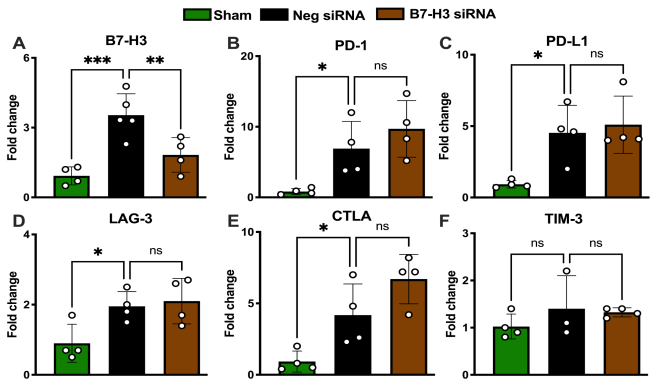


**Supplementary Fig 2:** Relative mRNA expression show significant increase of several immune checkpoints (B7-H3, LAG-3, PD-1, CTLA-4, TIM-3 and PD-L1) in the brain after stroke at (24 h of reperfusion) acute ischemic stroke. B7-H3 inhibition after stroke does not affect the induction of other immune checkpoints. (N = 4-5 per group) Statistical analysis was performed using a one-way ANOVA, followed by Dunnett's post hoc test.

**Additional motor functional assessment is less conclusive**

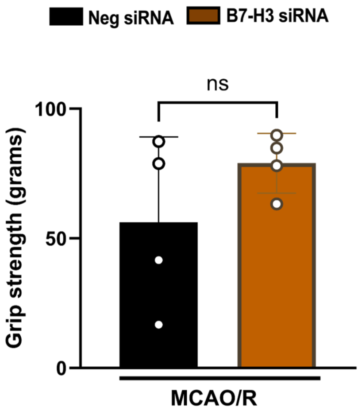


**Supplementary Fig 3:** Grip strength of MCAO/R mice following post-treatment of Neg-siRNA and B7-H3 siRNA at 7 days. Forelimb grip strength was assessed 7 days after MCAO/R in Neg-siRNA or B7-H3 siRNA. Data represented as mean ± SEM (N = 4-5 per group), Mann-Whitney test.
